# Aging is associated with a degeneration of noradrenergic-, but not dopaminergic-neurons, in the short-lived killifish *Nothobranchius furzeri*

**DOI:** 10.1101/2021.03.31.437850

**Authors:** Sara Bagnoli, Baldassare Fronte, Carlo Bibbiani, Eva Terzibasi Tozzini, Alessandro Cellerino

**Affiliations:** Laboratory of Biology (BIO@SNS), Scuola Normale Superiore, Pisa, Italy; Department of Veterinary Sciences, University of Pisa, Italy; Biology and Evolution of Marine Organisms Dep. (BEOM), Stazione Zoologica Anton Dohrn, Napoli, Italy; Fritz Lipmann Institute for Age Research, Leibniz Institute, Jena, Germany

**Author notes:** **Correspondence:** Alessandro Cellerino.

**Keywords:** Parkinson’s disease, killifish, *locus coeruleus*, neurodegeneration, alpha-Synuclein, protein misfolding, age-related disease, aging, animal model, telost

## Abstract

Parkinson’s disease (PD) is characterized by phosphorylation and aggregation of the protein α-Synuclein and ensuing neuronal death progressing from the noradrenergic *locus coeruleus* to midbrain dopaminergic neurons. In 2019, Matsui and colleagues reported in *Cell Reports* a spontaneous age-dependent degeneration of dopaminergic neurons and an even greater neurodegeneration of the noradrenergic neurons in the short-lived killifish *Nothobranchius furzeri*.

Given the great possible relevance of a spontaneous model for PD, we set to confirm their results by whole-brain clarification and 3D nuclei reconstruction to quantify total cell numbers. We observed an age dependent neurodegeneration limited to the *locus coeruleus* and not involving the *posterior tuberculum*. In addition, we observed the presence of phospho-Synuclein in the soma of *locus coeruleus* neurons detectable already at a young age and increasing during aging.

Our result demonstrates that *N. furzeri* models the early stages of PD, but not the degeneration of dopaminergic neurons.

## Introduction

Parkinson’s disease (PD) is one of the most prevalent neurodegenerative diseases characterized by a plethora of motor and non-motor symptoms due to dopaminergic and noradrenergic neuron loss. It has been long hypothesized that the pathogenic mechanism of neurodegeneration lies in the misfolding and subsequent aggregation of the neuronal protein α-Synuclein (α-Syn), leading to the formation of intracellular Lewy bodies (LB) and dystrophic Lewy neurites (LN) (Spillantini et al., 1997, Braak et al., 1999, Braak et al., 2002, Anderson et al., 2006, Samuel at al., 2016). Aggregates are associated with post-translational modifications of α-Synuclein and in particular phosphorylation of Serine 129 (pS129, Anderson et al., 2006). Aggregation and phosphorylation appear with a precise spatio-temporal progression that defines a staging for the pathological progression (Braak et al., 2002, Braak et al., 2003, Del Tredici and Braak 2012): the aggregation in its very early stages is observed in distal parts of the CNS and in particular the dorsal vagal nuclei in the brain stem then expands anteriorly hitting first the *locus coeruleus* (LC), subsequently the *substantia Nigra* (SN) *pars compacta* causing the onset of the typical motor symptoms and ultimately affecting the most anterior parts of the cortex. Propagation of α-Syn aggregates along the vagus nerve was recently demonstrated in animal models (Kim et al., 2019). *Locus coeruleus* and *substantia nigra* (SN) are the most studied nuclei in the context of PD. LC is affected at very early stages of PD and the extent of neuron loss is larger than in the SN, which is only affected during middle stages of the pathology (Braak et al., 2003, Zarow et al., 2003). It is also worth noting that presence of p-Syn immunoreactivity and LB are not an exclusive feature of PD. They are found in ~10% of clinically unimpaired subjects over the age of 60 with a distribution similar to PD, but showing lesser density of inclusions and a lesser degree of Tyrosine Hydroxylase (TH) reduction, as compared to what observed in PD, suggesting that p-Syn immunoreactivity and LB formation are already detectable at preclinical stages, before the onset of motor symptoms (Forno et al., 1969, Fumimura et al., 2007, Dickson et al., 2008).

Several approaches were undertaken to model PD in animal models. Dopaminergic loss can be induced in rodents by acute pharmacological treatments (for a reviews see Meredith et al., 2011, Johnson and Bobrovskaya, 2015), but this approach fails to model the progressive and age-associated nature of PD. An alternative approach envisages the genetic manipulation of either α-Syn or other PD-associated genes like LRRK2, PINK1, Parkin, or DJ-1 (Jagmag et al., 2016). However, the majority of PD cases do not have a clear genetic origin but are considered idiopathic (for a review on PD model limitations see Potashkin et al., 2010). Moreover, in most cases, the disease is modelled in young or, at best, in young adult animals. For a more precise modelling of the disease, it would be crucial to carry such studies in the context of old animals which is the natural environment for neurodegenerative diseases. Teleost fishes like Medaka and Zebrafish have also been utilized to model PD with similar approaches (Matsui et al., 2009, Flinn et al., 2013, Matsui et al., 2013, Uemura et al., 2015), thus showing the same limitations.

The teleost *Nothobranchius furzeri* (Killifish) is the vertebrate with the shortest captive lifespan and has emerged as a valuable experimental model for aging research. This species replicates the conserved aging hallmarks of mammals and in particular shows age-dependent gliosis (Terzibasi 2012), activation of immune response and spontaneous protein aggregation in neurons (Kelmer et al., 2020), making it a particularly valuable model for brain aging.

In 2019, Matsui and colleagues described a spontaneous syndrome in *N. furzeri* that shares some aspects of PD (Matsui et al., 2019). In their work they observed an age-dependent Syn accumulation associated with the formation of Synuclein aggregates and age-dependent neuronal loss in the LC and in a diencephalic dopaminergic population (*posterior tuberculum*) that project to the subpallium and is considered a putative homolog of mammalian midbrain dopaminergic neurons (Kaslin and Panula, 2001; Rink and Wulliman, 2001).

Given the great possible significance of this report, we set to replicate their study, but came to partially discordant results that are described here.

## Results

In the present study, we analyzed individuals from the *N. furzeri* population MZCS-222 that belongs to the same genetic clade of the population MZCS-24 studied by Matsui and the population MZM-0410 we have used in our previous studies. This population was collected later than MZM-0410 and is therefore likely to be more genetically heterogeneous and closer to the wild population. The median lifespan of this strain in captivity is around 6 months (Zak and Reichard, 2020).

We set to quantify the total number of TH+ cells in the LC and in the *Posterior tuberculum* (hypothalamus) in young (5 weeks) and old (37 weeks) animals. The size of the brain increases during this period and the spatial distribution of cells may vary between animals of different ages (Fig. 1A and video S3). In order to visualize and count all TH+ cells in the region of interest, we clarified the brain using the Sca/eS procedure (Hama et al., 2015, Hama et al., 2016) and reconstructed all the different TH+ nuclei of the *N. furzeri* in their entire 3D extent by scanning through the transparent brain (Fig. S1, S2).

**Fig.1.**
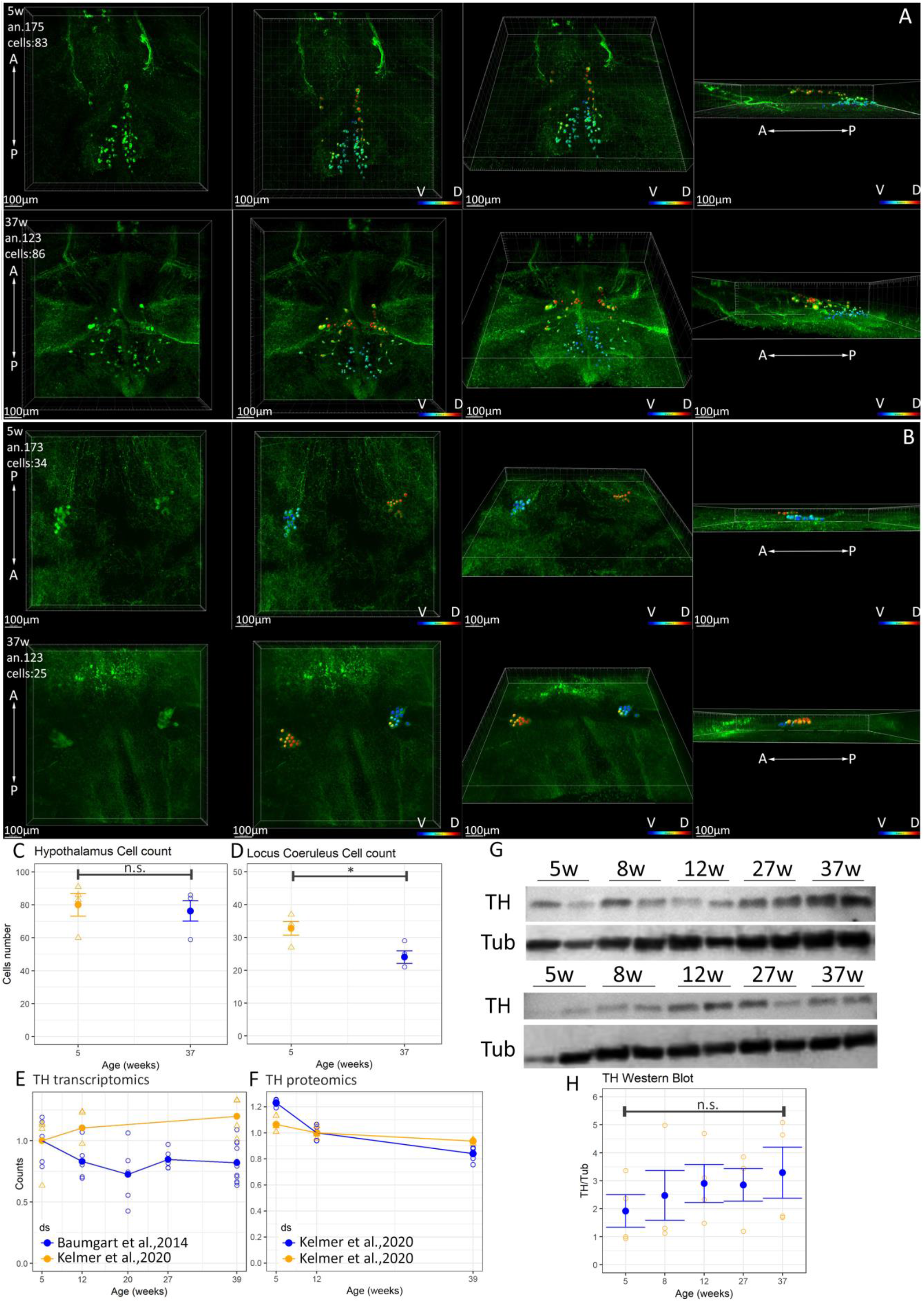
Assessment of neurodegeneration in LC and p*osterior tuberculum* and quantification of TH expression during ageing. A) Representative images of 3D reconstruction and cell counting in the *posterior tuberculum* nuclei of a young (5 weeks old) and old (37 weeks old) *N. furzeri.* V=ventral, D=dorsal, A=anterior, P=posterior. The other animals are reported in Fig. S2. The color code indicates the depth of the cells in the reconstructed volume. B) Representative images of a 3D reconstruction and cell counting of *locus coeruleus* of a young (5 weeks old) and old (37 weeks old) *N. furzeri.* V=ventral, D=dorsal, A=anterior, P=posterior. The color code indicates the depth of the cells in the reconstructed volume. The other animals are reported in Fig. S5. C) cell counts of *posterior tuberculum* nuclei in young and old animals. Each open dot represents a single animal and the solid dot the mean of the group and the error bar the SEM. D) cell counts of *locus coeruleus* nuclei in young and old animals. Statistical significant difference was assessed by Student’s t-test, n.s. indicates p > 0.05, * indicates p > 0.05. E) Quantification of TH mRNA expression. Two independent RNA-seq datasets (from Baumgart et al., 2014, and Kelmer et al., 2020) were analyzed. F) Analysis of TH protein expression by mass spectrometry. We assessed two independent proteomic datasets (both from Kelmer et al., 2020). Each open dot represents a single animal and the solid dot the mean of the group and the error bar the SEM. To assess the statistical significance of age-dependent regulation, we calculated Spearman correlation with age. In both cases p > 0.05. G) Images of TH protein expression by Western blot, complete blots are reported in Fig. S7. A total of four animals per each age were analyzed. H) Quantification of TH Western blot. The intensity of TH band was normalized with respect to the tubulin band. To assess statistical significance, we calculated Pearson’s correlation coefficient of the expression with age and respective pValue, the data are expressed as mean ±SEM.

The *N. furzeri posterior tuberculum* contains two clearly separated populations of TH+ cells (video S3), one comprised of smaller and more posteriorly- and ventrally-located cells and one comprised of larger and more anteriorly- and dorsally-located cells. These two nuclei are very similar to the nuclei 13 and 12 described in the zebrafish *posterior tuberculum* using TH immunofluorescence (Sallinen et al., 2009) and are proposed as homologous to the A-9 A-10 mammalian dopaminergic cluster comprising the *substantia nigra* (Kaslin and Panula, 2001; Rink and Wulliman 2001). We counted all cells in these two populations and could not detect a significant difference in their numbers between young and old animals (Fig. 1C), as opposed to the remarkable ~ 50% reduction reported by Matsui et al. Notably, the number of TH+ cells we detected was larger than the number reported by Matsui et al.: average 77 (range 59-86) vs. ~ 30. The complete tiles of optical sections and the 3D reconstruction of each individual are available to independently assess the correctness of our counting (Fig. S5).

As opposed to hypothalamic nuclei, a modest but significant decrease in the number of LC neurons (Fig. 1B, D video S4) was detected. Also in this case, the number of labelled cells we detected is larger than what reported by Matsui et al.: 33 (range 37-27) vs. ~ 20 and the amplitude of the cell loss much smaller (27% vs ~ 75%). Pictures representative of the 3D reconstructions of all the animals analyzed in the present study are presented in Figure S5.

Matsui et al. also reported a down-regulation of TH protein and transcript in the whole brain by Western blot and qPCR. We interrogated two independent public databases of RNA-seq (Fig. 1 E) and two independent databases of mass-spectrometry based proteomics of *N. furzeri* brain aging (Fig. 1 F) and we could not find any support for a down-regulation of TH neither at the transcript- nor at the protein-level. We repeated Western blot analysis and also in this case we could not detect a down-regulation of TH (Fig. 1G,H)

To assess pathological post-translational modifications of α-Syn, we used an antibody directed against phosphorylated S129 (pS129) in double-labelling with TH. Labelling of *posterior tuberculum* neurons did not reveal any somatic staining for pS129 in either young (5 weeks) or old (37 weeks) samples (Figure S6A). On the contrary, a punctate labelling of pS129 was readily detected in the LC neurons from old fish (Fig.2B). Remarkably, LC TH+ neurons contained punctate labelling of pS129 already at young age (Fig.2A). The only other cells showing early somatic labelling in the brain were TH+ cells in the vagal nuclei (Figure S6B). Quantitative analysis of the staining revealed that size and density of pS129+ puncta increased between young and old animals (Fig.2C-E). It should be noted that the cytoplasm of LC neurons in old animals shows a clear vacuolization, apparent as spherules devoid of TH immunoreactivity (Fig. 2Bb). Interestingly, pS129+ puncta are not present in the vacuoles but seem to be associated with- of in proximity of- their external surface (Fig.2Bb).

**Fig.2.**
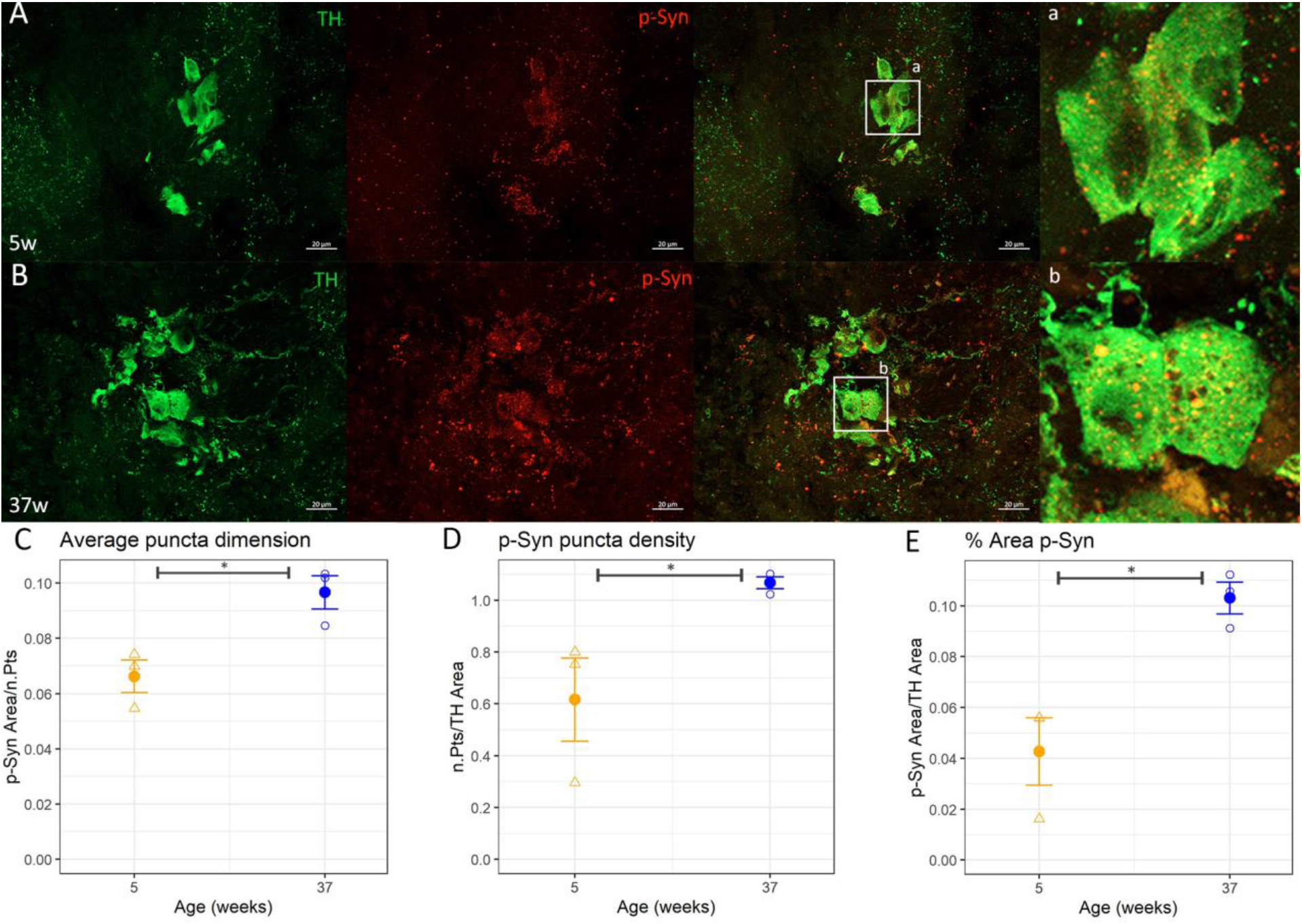
Double immunofluorescence for TH (green) and phospho-Synuclein (red) in *locus coeruleus* cells. A diffuse, punctate staining is visible in both young (5w, panel A,a) and old (37w, panel B,b) animals. Please note in the magnification shown in panel b the presence of cytoplasmic vacuoles. The dimension (panel C), the density (panel D) and the area (Panel E) of the puncta increases significantly with age. To test significance two tailed t-test was used * = p < 0.05, the data are expressed as mean ±SEM. Scale bars correspond to 20 μm.

We also assessed a general increase of pS129 staining in the whole brain and in the central telencephalon specifically (Fig. 3A,B), this region was chosen because age-dependent appearance of protein aggregates was previously demonstrated by us (Kelmer et al., 2020). Labelling outside of the LC was observed mostly in the form of labelled processes and, unlike in the LC and vagal nuclei, clearly labelled somata with punctuate labelling could not be detected. Both the total brain area covered by pS129 labelling and the density of labelling in the telencephalon were increased in old animals. Remarkably, pS129 in the telencephalon appeared to be associated, albeit not exclusively, to TH+ positive axonal fibers (Fig.3E). These dystrophic axons may originate from the LC, but we have no definitive proof of their identity.

**Fig.3.**
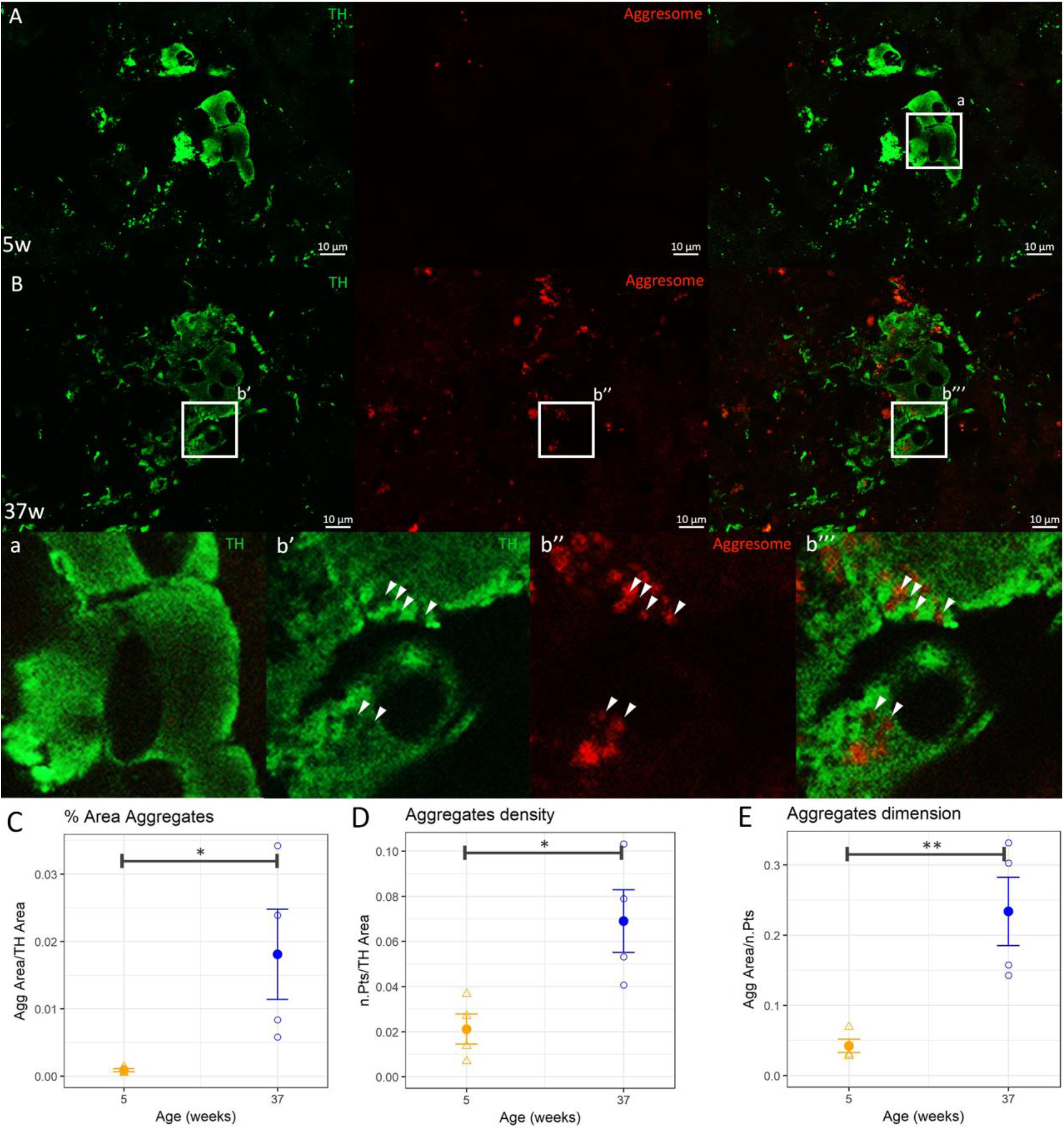
Double immunofluorescence for TH (green) and Aggregates (Proteostat Aggresome Dye, red) in *locus coeruleus* cells. Aggresome staining is absent in young animals (5w, panel A) and becomes visible in old animals (37w, panel B). In particular, aggregates are mainly located in vacuolized areas of cytoplasm devoid of TH staining (panels, b’, b’’, b’’’) that are not present in young animals (panel a). The area (panel C), the number (panel D) and the dimension (panel E) of the aggregates all increased significantly with ageing. To test significance two tailed t-test was used, * = p<0.05 and ** = p<0.01, the data are expressed as mean ±SEM. Scale bars correspond to 10 μm.

In addition, we investigated the presence of protein aggregates (aggresomes) in LC neurons from old fish. The Proteostat dye revealed a punctate staining in TH+ neurons from old, but not young fish (Fig. 4A,B). This difference was apparent and was also confirmed by quantitative analysis (Fig.4C-E). The localization of aggresomes in LC neurons was distinct from that of pS129 puncta, as protein aggregates were clearly contained within the cytoplasmic vacuoles (Fig. 3Bb’-b’’’).

**Fig.4.**
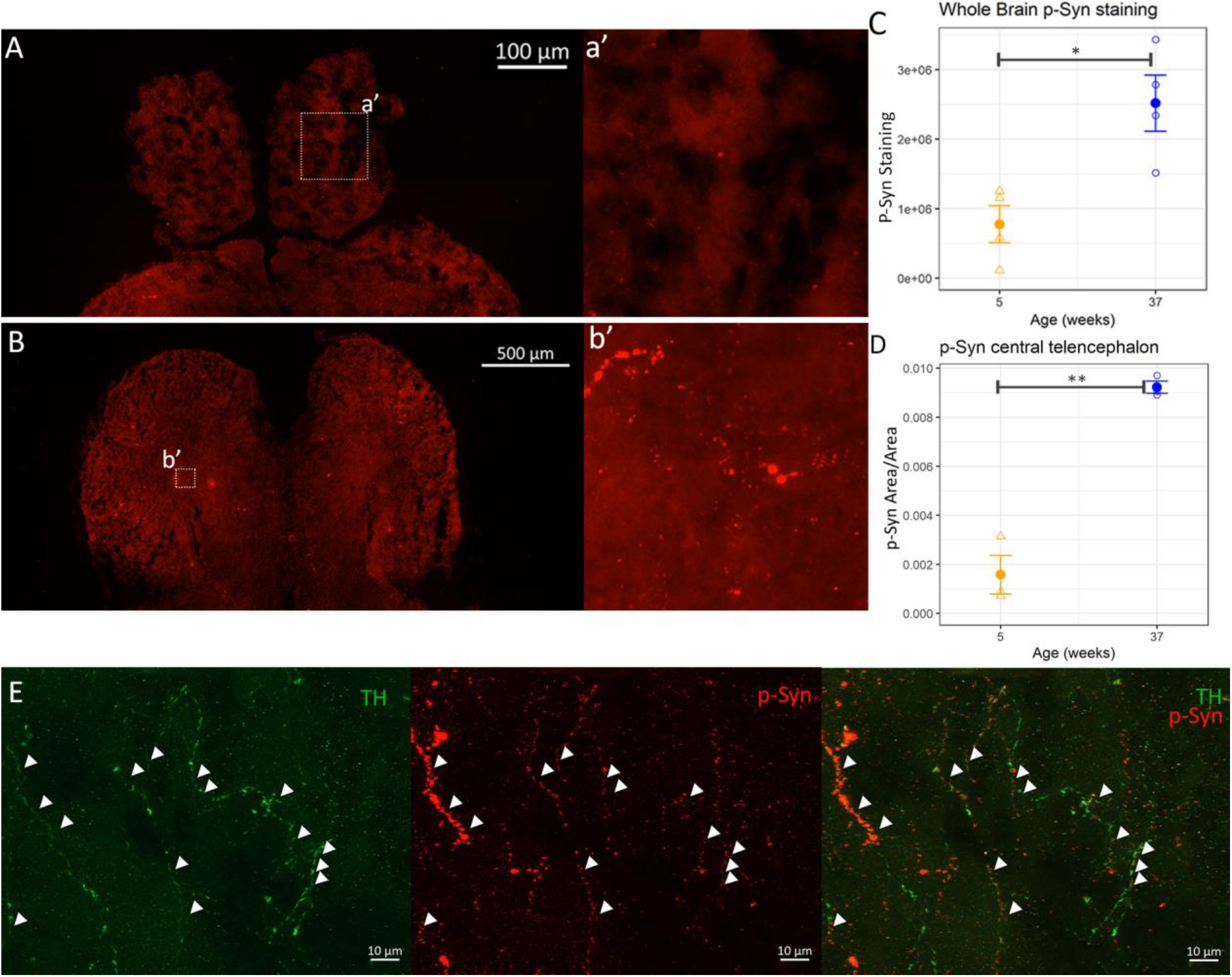
Phospho-Synuclein staining in other brain areas. We quantified the amount of phospho-Synuclein staining in the whole brain of young (5w panel A) and old (37w panel B) animals and we found a significant increase with ageing (panel C). We also focalized the analysis in the central telencephalon (panels a’, b’). Also in this area we found a statistical significant increase of phospho-Synuclein staining (panel D). To test significance two tailed t-test was used, * = p<0.05 and ** = p<0.01, the data are expressed as mean ±SEM. Apart from *locus coeruleus* and vagal cells, the staining of phospho-Synuclein is localized in processes, not exclusively colocalizing with TH positive fibers. We were able to find such staining associated with TH positive processes especially in old animals (37w) telencephalic areas (panel E).

## Discussion

In our paper, we report an age-dependent degeneration of noradrenergic neurons in the LC, but not of dopaminergic neurons in the *posterior tuberculum*, a brain region supposed to be the homolog of mammalian A8 and A9 dopaminergic populations. The results of our analysis contrasts with the results reported by Matsui et al. Two main factors may account for this discrepancy. In the first place, we used a different counting method and instead of counting cells in thick sections we clarified the entire brain and reconstructed the nuclei of interest in their entire 3D extent. Since *N. furzeri* brain grows considerably during adult life, it may be possible that a sampling protocol that identifies all cells in young animals may undersample the same cells in older animals. Indeed, our 3D reconstructions show that dopaminergic neurons of the posterior tuberculum are more widespread in brains from old fish. A second possibility is the existence of genetic differences between the two strains. Even if the population MZCS-222 we analyzed here and the population MZCS-24 analyzed by Matsui et al. are part of the same genetic clades and the distance between the two collection points is around 50 km, it cannot be formally excluded that the subtle genetic differences between the two stocks may have an impact on dopaminergic neuron maintenance. Lacking access to the population MZCS-24, we cannot exclude this possibility. Yet, our results exclude that degeneration of dopaminergic neurons is a general trait of wild-type *N. furzeri*.

Similarly, we do not find any evidence for a down-regulation of TH expression. Matsui et al. showed data from pooled animals, in our study we report the results of individual animals and they are supported also by non-targeted approaches such as RNA-seq and Mass-spectrometry based proteomics. This finding is not surprising, giving the presence of a large number of TH+ cells in other brain regions, such as the olfactory bulb (Fig. S1). A specific loss of LC and *posterior tuberculum* neurons would have a negligible impact on the abundance of TH in the whole brain.

Neurodegeneration is associated with accumulation of the pathogenic post-translational modification pS129. The condition we describe, with accumulation of pS129 is reminiscent of the prodromic, pre-symptomatic stages of PD as described by (Forno et al., 1969, Braak et al., 2003, Dickson et al., 2008) that show staining with pS129 in the vagal nuclei and LC that precedes the degeneration of dopaminergic neurons and the onset of motor symptoms. Recently, it has been proposed that liquid-liquid phase separation could be the initial mechanism of α-Syn nucleation, and it has been demonstrated *in vitro* how a phosphomimetic mutation (S129E) increases the rate of liquid-droplet formation and reduces the critical concentration of protein required for the beginning of the phase separation (Ray et al., 2020), in accordance with the notion that phosphorylation of Serine 129 site is critical for aggregation of α-Syn (Samuel et al., 2016).

It should be noted that we do not have evidence for formation of Lewy bodies or large aggregates. The labelling of pS129 remains indeed punctate and does not overlap with staining for aggresome that is localized within cytoplasmic vacuoles. This labelling may therefore correspond to an initial conformation change that precedes the formation of aggregates.

In summary, we describe a condition that is reminiscent of the presymptomatic stage of PD. *N. furzeri* could then be used in the future to identify modifying factors that are responsible for the pathological transition.

## Acknowledgements

We thank Martin Reichar for providing us eggs of the MZCS-222 strain. The work was partially supported by the grant SNS2019 to AC.

## Author Contributions

SB and AC designed the experiments, SB performed the experiments with help from ETT, BF and CB. AC and ETT supervised the work. SB and AC wrote the paper with help from all the authors.

## Star Methods

### LEAD CONTACT

Further information and requests for resources and reagents should be directed to and will be fulfilled by the lead contact, Alessandro Cellerino

### MATERIALS AVAILABILITY

Images (.tiff) and IMARIS 3D reconstructions are deposited and available at the following link: https://data.mendeley.com/datasets/4njyjn4s86/draft?a=9bb9bc24-ae3a-420b-ab68-8edd7facb8a0

### DATA AND CODE AVAILABILITY STATEMENT

This study did not generate any unique datasets or code.

### EXPERIMENTAL MODEL AND SUBJECT DETAILS

#### Fish maintenance and sampling

MZM-222 were hatched and housed locally in a Tecniplast Zebtech system with automatized water flow, pH and salinity control. All the animals were hatched, fed and maintained as described in detail in (Terzibasi et al., 2008).

The protocols of fish maintenance were carried out in accordance with all animal use practices approved by the Italian Ministry of Health (Number 96/2003a) and the local animal welfare committee of the University of Pisa.

Eggs were maintained on wet peat moss at room temperature in sealed Petri dishes. When embryos had developed, eggs were hatched by flushing the peat with tap water at 16–18 °C. Embryos were scooped with a cut plastic pipette and transferred to a clean vessel. Fry were fed with newly hatched *Artemia nauplii* for the first 2 weeks and then weaned with finely chopped *Chironomus* larvae. Starting at the fourth week of life, fish were moved to system tanks. The aquarium room’s temperature was set at a constant 27 °C. At the desired age, fish were sacrificed via anesthetic overdose (Tricaine, MS-222), in accordance with the prescription of the European (Directive 2010/63/UE) and Italian law (DL 26/04-03-2014), and the brain was immediately extracted under a stereomicroscope and fixed overnight in a solution of PFA 4% in PBS. Samples used for AbSca/e procedure were then gradually dehydrated by sequential steps of incubation in solutions with growing EtOH concentration (25%, 50%, 75%) and finally stored at −20° in EtOH 90% until use.

Samples used for immunofluorescence were instead incubated overnight in sucrose 30% at 4°C and the day after were included in cryo-embedding medium (Tissue-Tek^®^ O.C.T., Sakura Finetek). Serial slices of 25 μm of thickness were then cut using a Leica cryostat and collected on Superfrost plus slides^®^ (Thermo scientific).

### METHOD DETAILS

#### Sca/eS procedure

Sca/eS technique is a clarification technique (Hama et *al*., 2015, Hama et al., 2016).

We followed the principal steps of the AbSca/e technique, which is a subtype of Sca/eS thought especially to combine immunofluorescence and tissue clarification, adapting some of the steps to the considerably smaller size of *Nothobranchius furzeri* brains. All the steps and the solutions used for this procedure are summarized in tables 1 and 2.

**Table 1.** Composition of the solution used for the Sca/eS protocol. The composition is taken as indicated in Hama et al,. 2015

| Ingredients | Sca/eA2 | Sca/e B4(0) | Sca/eS0 | Sca/eS4 | AbSca/e | AbRinse |
| --- | --- | --- | --- | --- | --- | --- |
| D-(-)-sorbitol (w/v)% | - | - | 20 | 40 | - | - |
| Glycerol (w/v)% | 10 | - | 5 | 10 | - | - |
| Urea (M) | 4 | 8 | - | 4 | 0.33 | - |
| Triton-X-100 (w/v)% | 0.1 | - | - | 0.2 | 0.5 | 0.05 |
| Methyl-β-cyclodextrin (mM) | - | - | 1 | - | - | - |
| γ-Cyclodextrin (mM) | - | - | 1 | - | - | - |
| N-acetyl-L-hydroxyproline (w/v)% | - | - | 1 | - | - | - |
| DMSO (w/v)% | - | - | 3 | 25 | - | - |
| PBS | - | - | 1x | - | - | 0.1x |
| BSA (w/v)% | - | - | - | - | - | 2.5 |

**Table 2.**
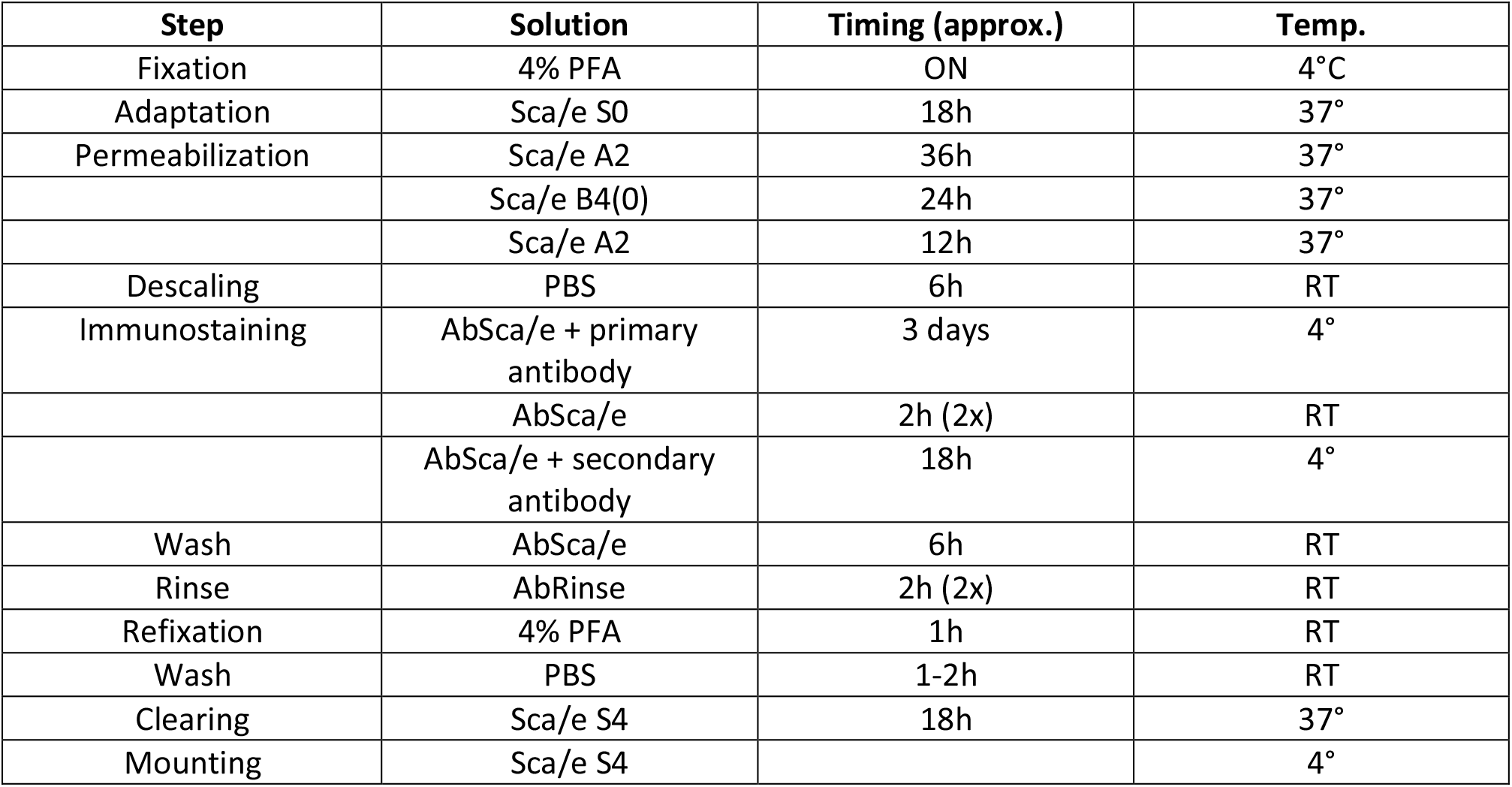
Sca/eS procedure. This is the procedure we adapted from Hama et al., 2015 for clearing and staining of *Nothobranchius furzeri* brains. In the table are reported the solution required for each step with the time and temperature of incubation. ON: over night, RT: room temperature.

Briefly, the samples were re-hydrated via incubation in solutions containing decreasing concentrations of ethanol (90%->75%->50%->25%->PBS) and adapted with the S0 solution for 18h at 37°C. Samples were then permeabilized with sequential incubations in A2-B4(0)-A2 solutions at 37°C. After permeabilization the samples were de-Sca/ed trough incubation in PBS for 6h at RT followed by incubation with primary antibody in AbSca/e solution for 3 days at 4°C, rinsed twice in AbSca/e for two hours each and incubation with secondary antibody for 18h at 4°C. The samples were then rinsed for 6h in AbSca/e and subsequently in AbRinse solution twice for 2 hours. After a re-fixation step in PFA4% for 1 h and a rinse in PBS for another hour the samples were finally clarified in Sca/e S4 for 18h at 37° and maintained in Sca/eS40 at 4° until imaging. The Sca/eS4 solution was also used as an embedding medium for imaging.

#### Immunofluorescence and Proteostat™ Aggresome staining

We performed immunofluorescence experiments on horizontal cryo-sections of 25μm thickness and proceeded as previously described (Tozzini et al., 2012). Briefly we washed the sections in PBS to remove the cryo-embedding medium; then we performed an acid antigen retrieval step (Tri-sodium citrate dehydrate 10mM, tween 0,05%, pH6). Afterwards, we stained the section with Aggresome (ProteoStat™ Aggresome Detection Kit, Enzo Life Sciences Inc.; for more details see Shen et al. 2011) as follows: we applied a solution 1:2000 of Aggresome dye in PBS for 3 minutes, rinsed in PBS and left the sections immersed in 1% acetic acid 40 minutes for de-staining. We applied blocking solution (5%BSA, 0,3% Triton-X in PBS) for 2 hours, then the primary antibody at proper dilution in a solution of 1% BSA, 0,1% triton in PBS, and incubated the samples overnight at 4°C. After rinsing in PBS, the following day secondary antibody was applied at a 1:400 dilution in the same solution used for the primary antibody. After 2 hours, slides were rinsed 3 times with PBS and mounted with a specific mounting medium added with nuclear staining (Fluoroshield DAPI mounting, Sigma-Aldrich). The antibodies utilized were: Thyroxine hydroxylase (Rabbit Monoclonal, ab75875, Abcam), anti-α-Synuclein Phospho (Ser129) (Mouse Monoclonal, 825702, Biolegend), Alexa Fluor 568 Goat anti Mouse (A11004) and Alexa Fluor 488 Goat anti Rabbit (A11008) (Invitrogen).

#### AbSca/e samples images acquisition and processing

To analyze the AbSca/e treated samples we acquired images on a Leica Ire2 confocal microscope using a 20× objective, exciting with an Ar-laser (488 nm).

We acquired 1024×1024 pixel sequential focal planes along the z axis, at a distance of 1.5 μm each, of the regions of interest and then realized and processed the 3D reconstruction utilizing the Bitplane Imaris™ software. First, we applied median filtering to reduce background, then we adjusted the tone curve accordingly to the characteristics of the imaged sample.

To count the TH+ cells in the *Locus coeruleus* and the hypothalamus of *Nothobranchius furzeri* brain we utilized the Imaris™ ‘Ortho Slicer’ function to optically isolate portions of the z-stack and the function ‘Spots’ to manually count of every TH positive cell.

We analyzed a total of 4 animals per age.

#### Immunofluorescence images acquisition and processing

Images were acquired using a Zeiss Axiovision microscope equipped with Apotome slide and subsequently processed using the suite Zen Blue.

Single area images (*Locus coeruleus,* hypothalamus, vagal cells, telencephalon) were acquired using two different magnifications (objectives 40× and 63× both oil immersion) as z-stacks formed by multiple focal planes at a distance of 0.5μm each.

### QUANTIFICATION AND STATISTICAL ANALYSIS

#### Whole brain p-Syn quantification

To perform a statistical quantification of p-Syn staining in whole *N.furzeri* brain, we cut 25μm thick cryo-slices in series of three, processing only one whole series per animal (i.e. 25μm every 75μm of tissue) for p-Syn and TH staining as described in the *Immunofluorescence and Proteostat™ Aggresome staining* section. We analyzed four animal per age (5w and 37w). We then acquired tiles at 20× using an AxioScan Zeiss microscope to obtain images of whole sections, using constant exposure time and applying the ‘Best Fit’ algorithm in ZenBlue.

The whole section images were then exported as .tif files and analyzed using Adobe Photoshop as described as follows:

Three copies of the image were created;

– to the first, a low threshold (value 15) was applied to isolate the area of the section, the area was then selected (Selection -> Color interval) and analyzed (Image -> Analysis -> Record Measurements); finally the summary containing the area (expressed in pixels) was exported.
– to the second copy a blur filter was applied (Filter -> Blur Gallery -> Field Blur, applied 80px level), and this image was used as a first background subtraction method.
– to the third copy the blurred one of the second step was applied (Image -> Apply Image -> select second copy with the fusion option subtract) to reduce the background. A threshold (value 40) was then applied to isolate the staining. Again, the staining was selected (Selection -> Color interval) and analyzed (Image -> Analysis -> Record Measurements) and the summary result containing the total px of staining was exported.

The area of each section and the area of the staining were registered for each animal in an Excel file and the percentage of stained area was calculated as follows (R=ratio, S=pixels of staining, A=pixels of section, i=section, n=total number of section per animal):

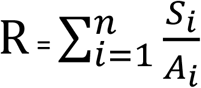

#### Central telencephalon p-Syn analysis

We used images collected with the same methodology used for the whole brain p-Syn quantification. We analyzed three animals per age (5w and 37w). We selected the four sections corresponding to the most central portion of the telencephalon along the dorso-ventral axis, and we isolated a square of 300px × 300px (A) into the central region on each section. The images were analyzed with the same methodology utilized for the whole sections and the percentage of stained area was calculated as follows (R=ratio, S=px of staining, A=area, i=section):

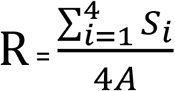

#### p-Syn and Aggresome analysis into the locus coeruleus

We analyzed images of *Locus coeruleus* acquired at a Zeiss AxioScan microscope with Apotome slide on a 40x magnification, all with the same exposure time and to which we applied the ‘Best Fit’ option in ZenBlue. We analyzed three animals per age for p-Syn and four animals per age for Aggresome. We acquired a z-stack of the area and then analyzed the single focal plane presenting the majority of TH staining. We used the open license software Icy (http://icy.bioimageanalysis.org/) for spot analysis. To the selected z-plane we applied the algorithm ‘Best Thresholder’ (method ‘Otsu’) on the TH channel to isolate the *Locus coeruleus* cells. Since TH is not distributed homogeneously into the cytoplasm but presents vacuolization containing aggregates, we adjusted manually the selected ROI to include areas containing aggregates. We then used the algorithm ‘Spot Detector’ imposing to perform the analysis only in the selected ROI with the following parameters:

Preprocessing: red channel
Detector: Detect bright spots over dark BG, scale 1 (for p-Syn) or scale 2 (for Aggresome) Region of interest: ROIFixedFromSequence
Output: export to ROI

The ROIs created (containing the data of both the area of TH and each single spot detected) were exported in an excel file and the following parameters were calculated:

– Percentage of stained area: spots area/ TH area
– Mean spots abundance: number of spots / TH area
– Mean spot dimension: spots area / number of spots

#### Western blot

We performed western blot experiments to quantify the expression of Thyroxine Hydroxylase in the brains of MZ-222 *Nothobranchius furzeri* at various ages. The brains were extracted, immediately stored at −80° and then homogenized for 30 seconds using GelD2 buffer with addition of protease and phosphatases inhibitors. After centrifugation for 10 minutes at 16000 rpm, supernatant was taken, quantified using the BCA kit (Thermofisher) and stored at −80° for further use.

To perform the western blot we run 20μg of protein extract derived from four samples per age, and run them in precast gels (AnyKD Mini-Protean TGX Gels, Biorad) for 35 minutes at 100V. Afterwards the samples were blotted on a nitrocellulose membrane for 35 minutes at 150V. The membranes were then imaged with a Chemidoc XRS scanner using the Quantity one Biorad software and band intensity quantification was performed using the open source ImageJ software. The two membranes were processed in parallel throughout the experiment and the acquisition phase. We analyzed four animals for each age (5,8,12,27,37 weeks).

#### RNA-seq and proteomics data analysis

To assess TH expression we realized graphics of RNA expression starting from RNA-seq data derived from public cross-sectional experiments (Baumgart et al., 2014, Kelmer et al., 2020). We combined two RNA-seq datasets of brain aging in *N. furzeri*: a dataset covering five time points (5, 12, 20, 27, and 39 weeks) and 5 biological replicates for each age and the second containing 4 replicates per ages: 5,12 and 39 weeks. These ages correspond to sexual maturity, young adult, adult (as defined by a decrease in growth rate), median lifespan and old (~30% survivorship)(Baumgart et al., 2014, Groth et al. 2014), five additional animals at 39 weeks (Reichwald et al., 2015, Petzold et al. 2015). In total, these represent 46 different samples spanning five different ages. We divided the counts of the two datasets for the average expression of their respective 5w samples to normalize their expression and to be able to combine the datasets.

We also analyzed publicly available proteomic data (Kelmer et al., 2020); we combined two datasets containing the first with five replicates for age (5, 12, 39 weeks) and the second with three replicates for age. The first dataset was a combination of two separate experiments performed utilizing tandem mass tag and analyzing the same 12w animals twice in two different contrasts: 39w vs 12w and 5w vs 12w., Therefore, we combined the data normalizing the protein expression dividing them for the mean value of the 12w animals. We applied the same normalization to the second dataset and plotted the resulting values.

For both transcriptomics and proteomics data we performed statistical analysis calculating the Spearman correlation value, the pValue adjusted and the FDR to assess the statistical significance of the observed expression variability.

All the analysis were performed utilizing the suite R.

#### KEY RESOURCES TABLE

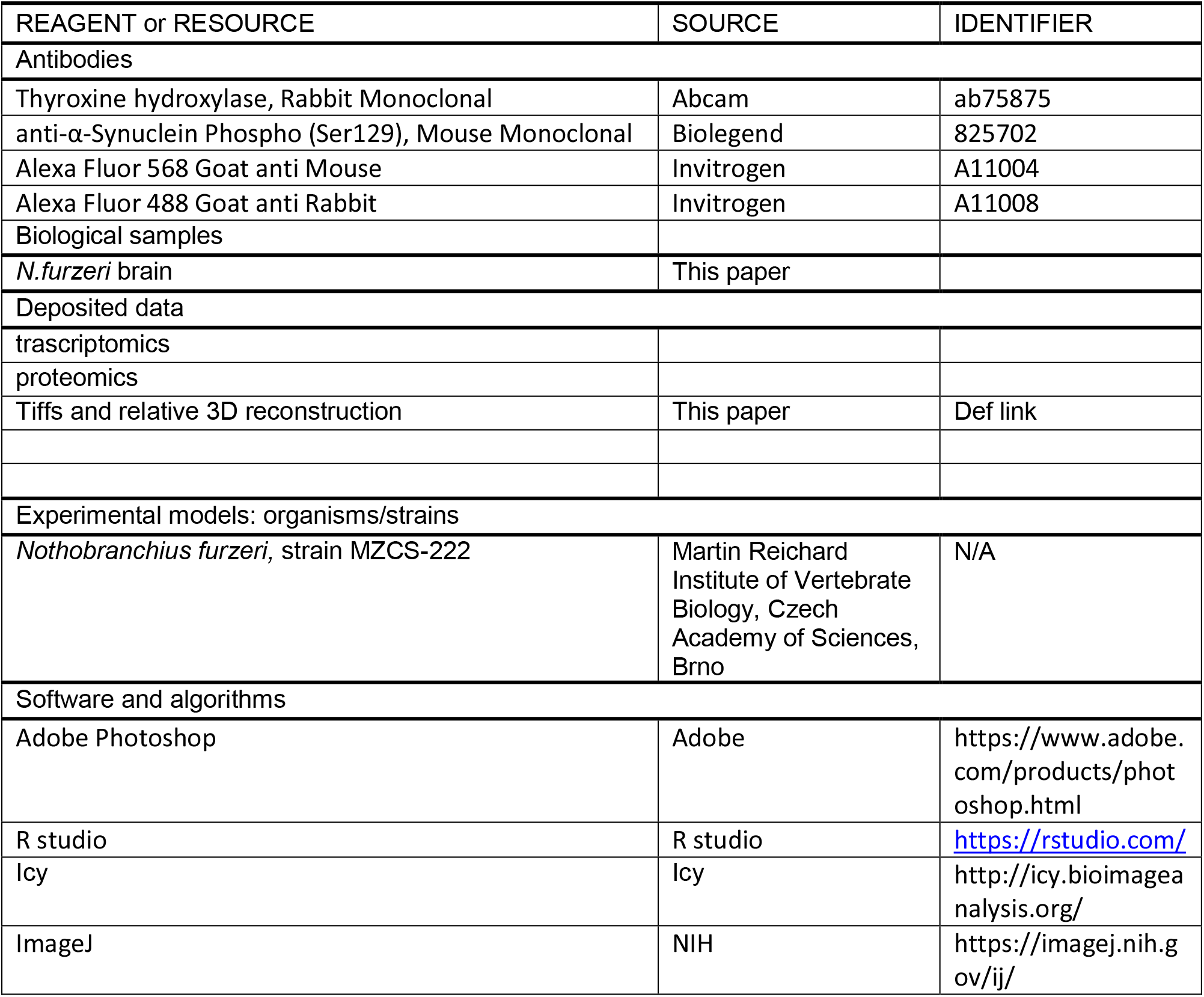
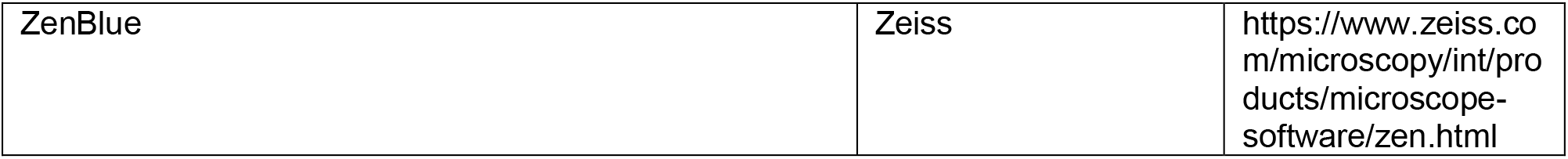

### Multimedia

Video S3. Comparison of 3D reconstruction of *locus coeruleus* of young (5w) and old (37w) *N.furzeri* with cell count.

Video S4. Comparison of 3D reconstruction of *posterior tuberculum* (hypothalamus) of young (5w) and old (37w) *N.furzeri* with cell count.

Link to complete tiff set and 3D IMARIS reconstructions:

