## Supplementary figures for "Aging is associated with a degeneration of noradrenergic-, but not dopaminergic-neurons, in the short-lived killifish *Nothobranchius furzeri*"

**Primary Supplemetal Information**

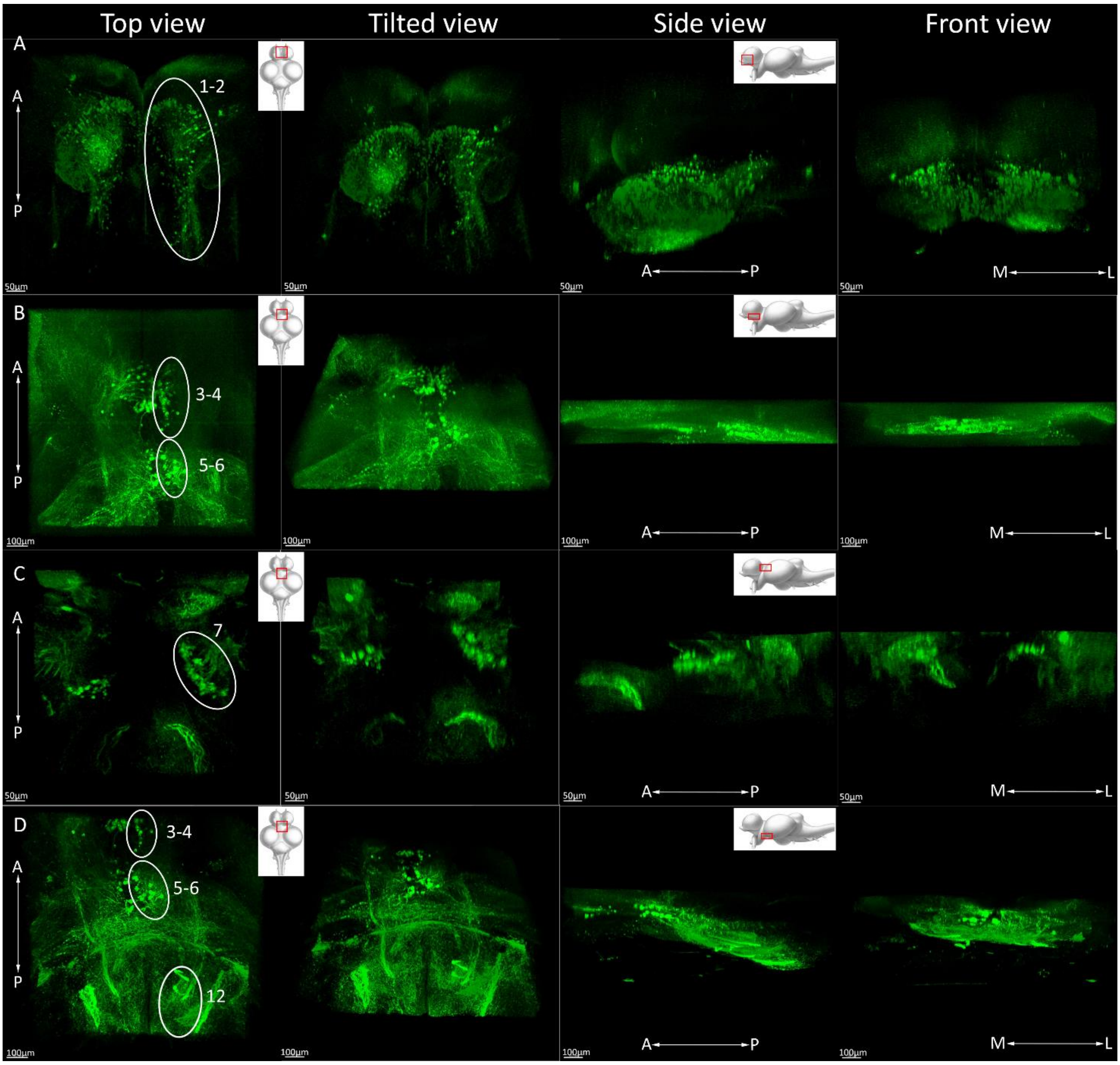


Fig.S1 Localization of the TH+ nuclei in the brain of *Nothobranchius furzeri*, anterior. As the main reference for nuclei identification a Zebrafish map from Sallinen et al. was used. Panel A: main nuclei of the olfactory bulbs and ventral telencephalon (homolog to nuclei 1-2 from Sallinen et al., 2009). Panel B: caudal telencephalic nuclei and rostral diencephalic nuclei (homolog to nuclei 3-4 and 5-6 from Sallinen et al., 2009). Panel C: periventricular pretectal nuclei (homolog to nuclei 7 from Sallinen et al., 2009). Panel D: caudal telencephalic nuclei, rostral diencephalic nuclei and beginning of periventricular organ, *Posterior tuberculum* (homolog to nuclei 3-4, 5-6 and 12 from Sallinen et al., 2009). A=anterior, P=posterior, M=medial, L=lateral.

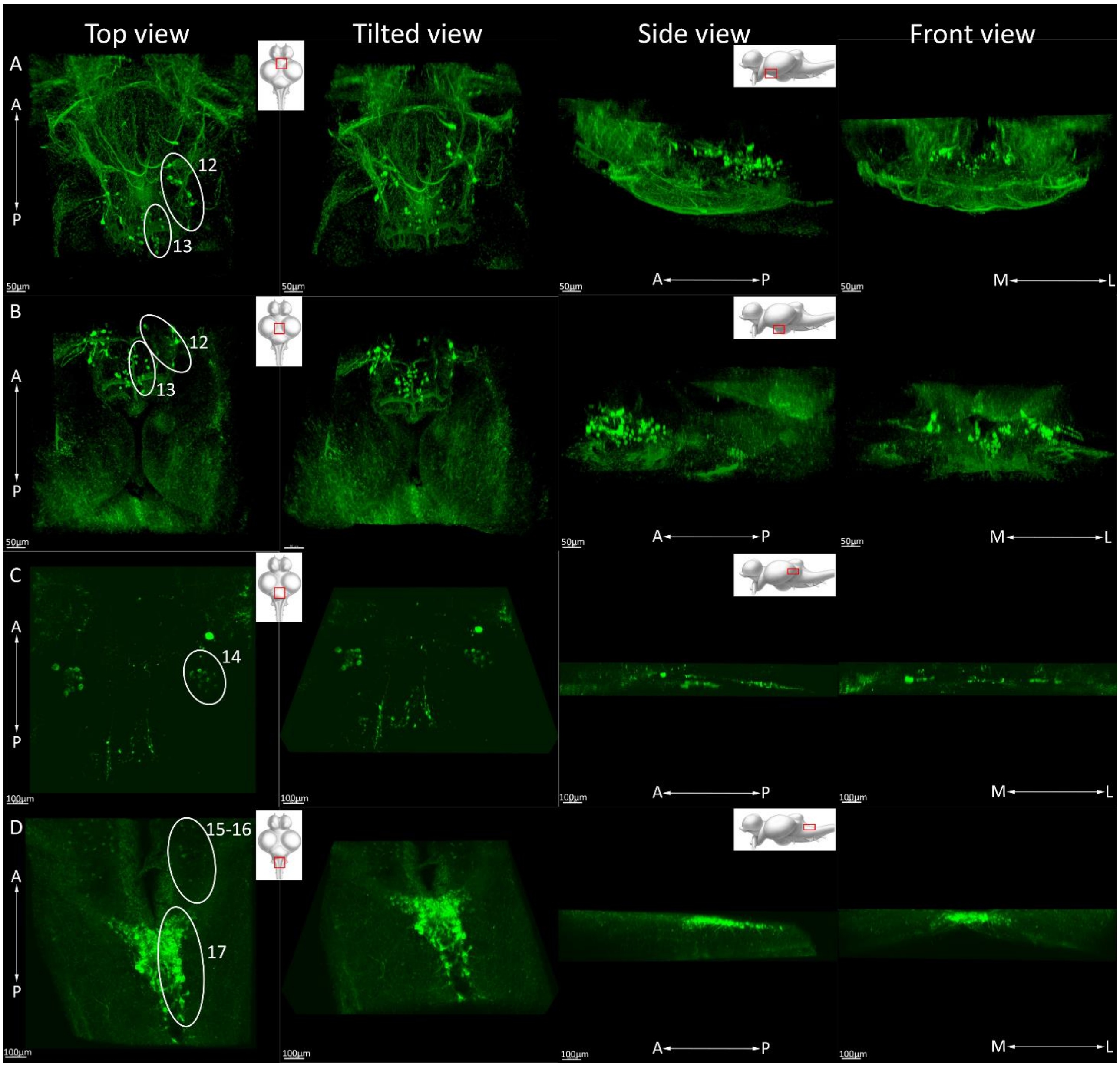


Fig.S2 Localization of the TH+ nuclei in the brain of *Nothobranchius furzeri*, posterior. As the main reference for nuclei identification a Zebrafish map from Sallinen et al. was used. Panel A: main diencephalic nuclei of the *Posterior tuberculum*: periventricular organ and periventricular hypothalamus (homolog to nuclei 12-13 from Sallinen et al., 2009). Panel B: more caudal view of main diencephalic nuclei of the *Posterior tuberculum*: periventricular organ and periventricular hypothalamus (homolog to nuclei 12-13 from Sallinen et al., 2009). Panel C: *Locus coeruleus* nuclei (homolog to nuclei 14 from Sallinen et al., 2009). Panel D: Vagal nuclei (homolog to nuclei 15-17 from Sallinen et al., 2009).

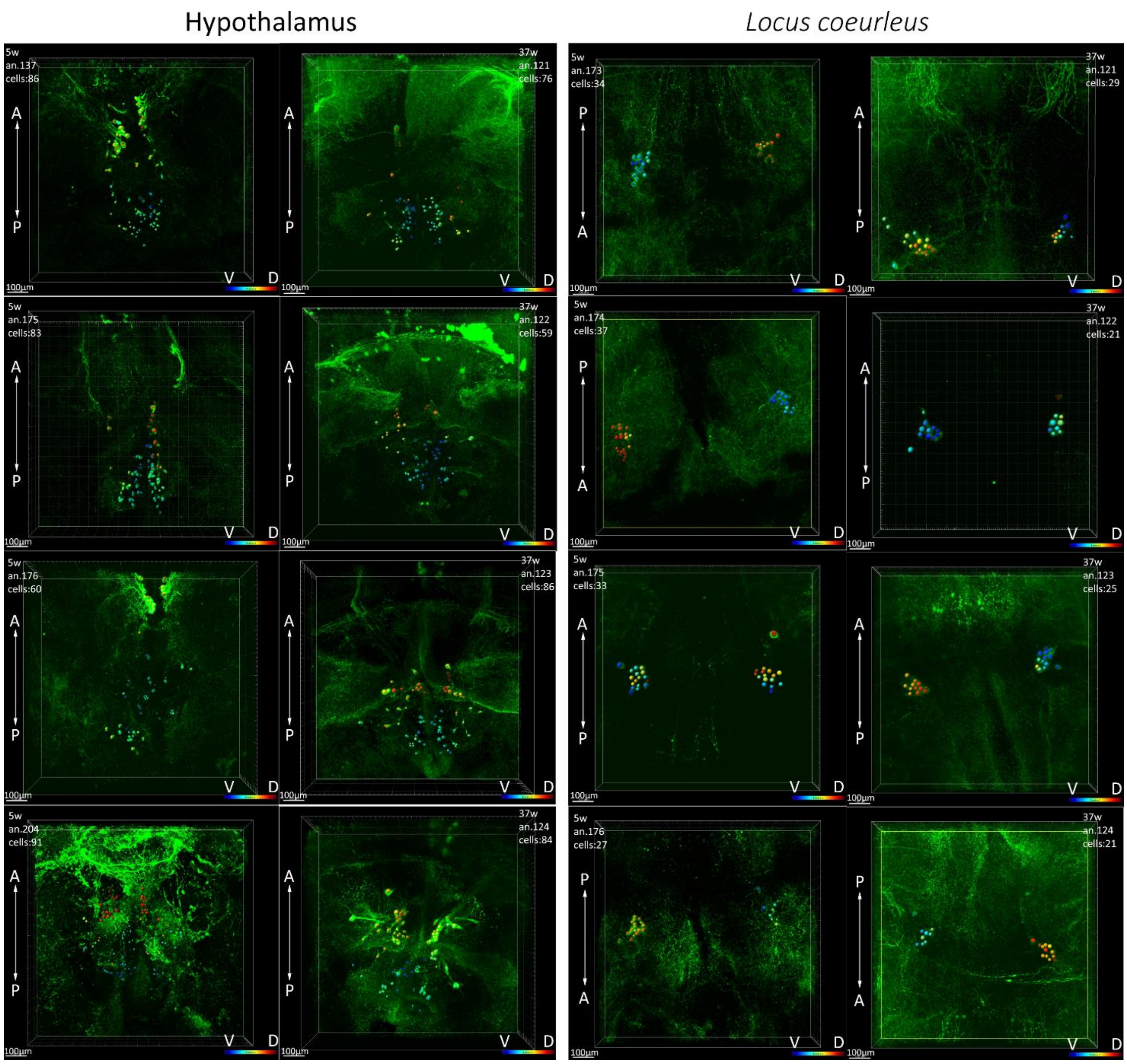


Fig.S5 Comparison of all 3D reconstructions with cell count of young (5w) and old (37w) *N.furzeri posterior tuberculum* (hypothalamus) and *locus coeruleus*. A= anterior, P=posterior, V=ventral, D=dorsal.

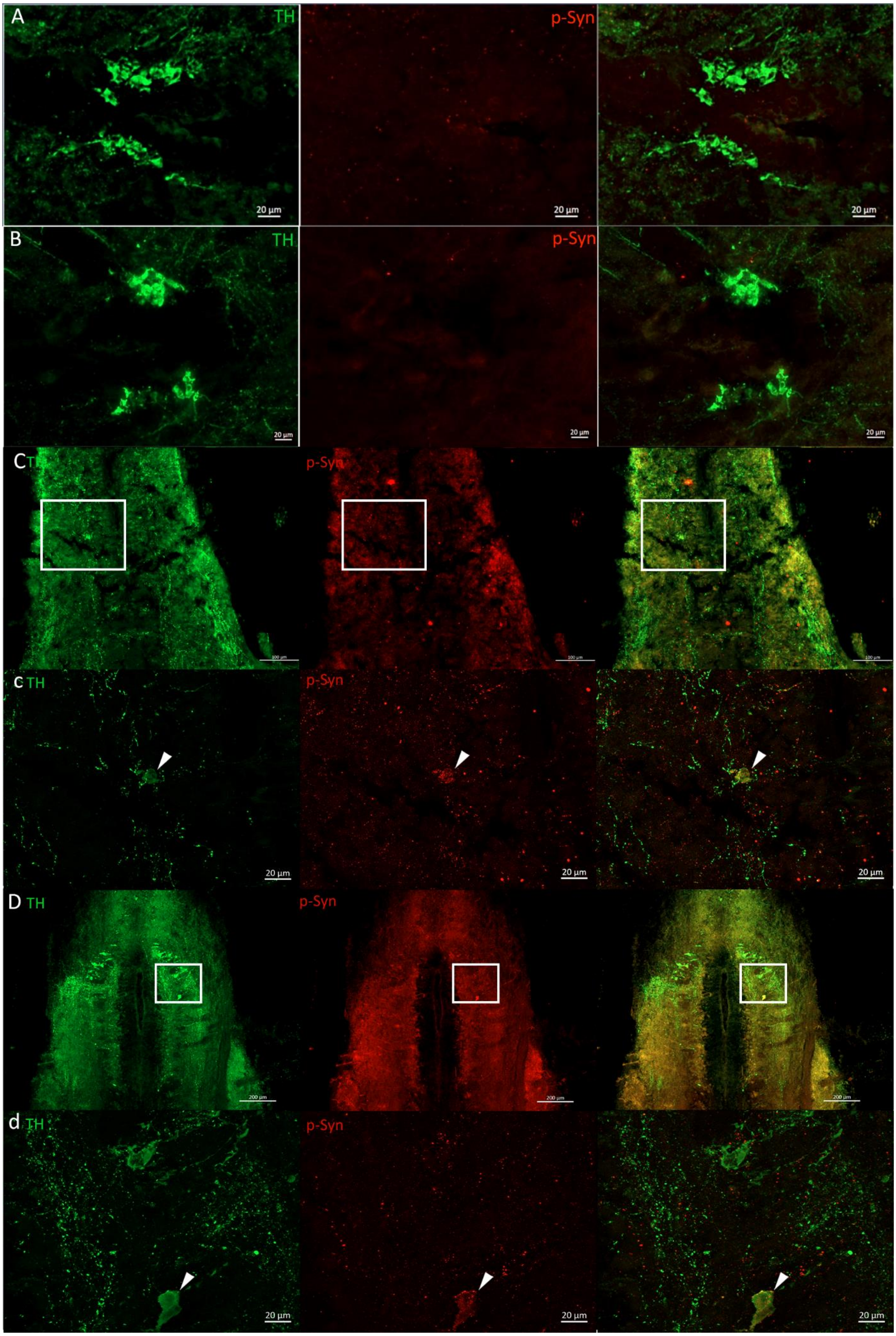
A
TH
p-Syn
20 µm
B
TH
p-Syn
20 µm
C
TH
p-Syn
c
TH
p-Syn
20 µm
D
TH
p-Syn
200 µm
d
TH
p-Syn
20 µm

Fig.S6 TH (green) and phospho-Synuclein (red) staining in *posterior tuberculum* (hypothalamus) and vagal nuclei. Phospho-Synuclein staining was absent from hypothalamic cell bodies both at young (5w, panel A) and old age (37w, panel B), while a somatic, punctate staining was present already at young age (5w, panels C,c) in cell bodies located in the vagal nuclei and remained evident also in old animals (37w, panels D,d). In panel C scale bar corresponds to 100 µm and in panel D to 200 µm.

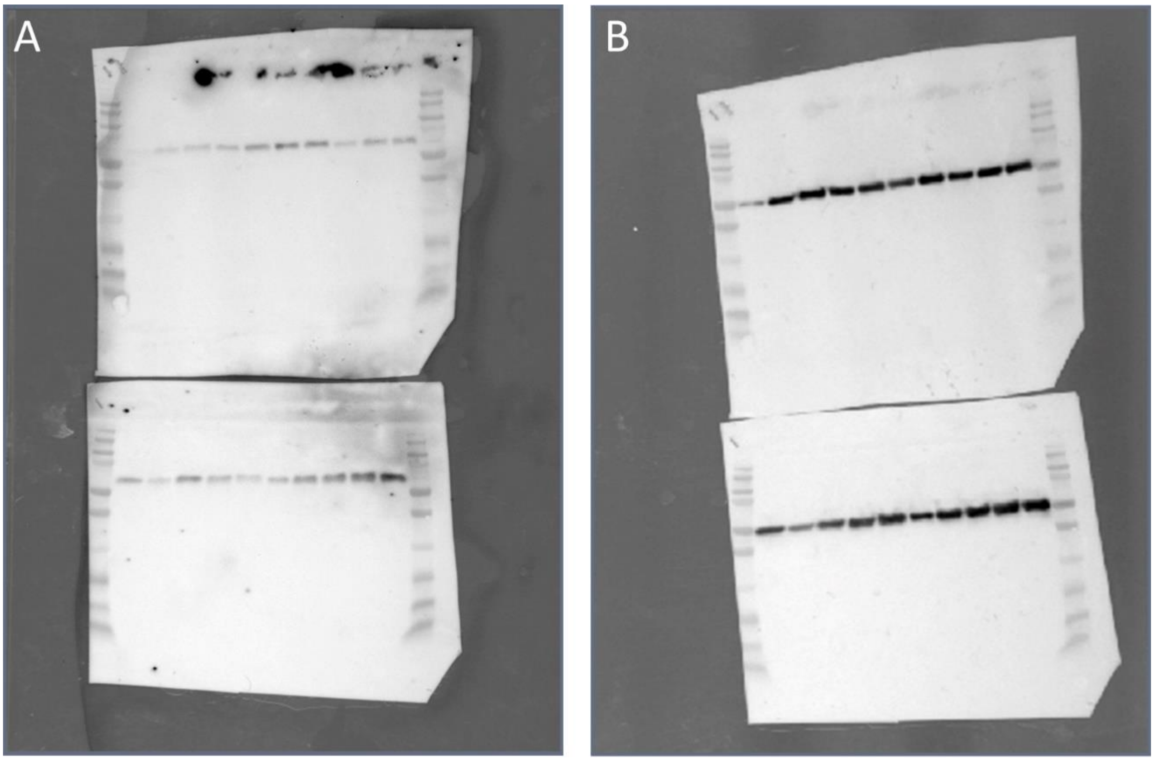


Fig.S7 Integral images for TH western blot. On panel A the integral original image for TH and on panel B the integral original image for Tubulin.
